# Exon Nomenclature and Classification of Transcripts (ENACT): Systematic framework to annotate exon attributes

**DOI:** 10.1101/2024.06.07.597685

**Authors:** Paras Verma, Deeksha Thakur, Shashi Bhushan Pandit

**Affiliations:** Indian Institute of Science Education and Research (IISER) – Mohali, Knowledge City, Sector-81, SAS Nagar, Manauli PO 140306, India

**Keywords:** Alternative splicing, isoform diversity, exon nomenclature, alternate transcription, and alternate translation

## Abstract

**Motivation:** Isoform diversity is known to enhance a gene’s functional repertoire. Despite studies on transcriptome diversifying processes (Alternate splicing/transcription), their extent and correlated impact on proteome diversity remains rudimentarily understood.

**Results:** The current study presents an innovative framework, “Exon Nomenclature and Annotation of Transcripts,” that centralizes exonic loci while integrating protein sequence *per entity* with tracking and assessing splice site variability. The resulting annotation from framework enables exon features to be tractable, facilitating a systematic analysis of isoform diversity. Our findings and case studies unveil systemic exon inclusion’s roles in regulating diversity in CDS region.

**Availability and implementation:** All data generated during this study are publicly available at www.iscbglab.in/enactdb/. Associated algorithmic procedures have been described in the methods section.

**Supplementary information:** PDF file enclosing supplementary data attached.

## Introduction

Gene architecture in eukaryotes facilitates generation of ≥ 1 mRNA per gene through a differential combination of exons. The exonic region determination and their subsequent combination is under regulation of several co-and post-transcriptional processes (Baralle and Giudice, 2017; Tapial, et al., 2017; Verta and Jacobs, 2022). Among these alternative splicing (AS) is widely studied, followed by alternative transcription initiation (ATRI) and termination (ATRT) (Kamieniarz-Gdula and Proudfoot, 2019; Ni, et al., 2010). Additional mechanisms such as Alternate translation initiation (ATLI) and termination (ATLT) acting at translation level also contribute to proteome diversity, where these processes are widely associated with leaky scanning, re-initiation, inclusion of upstream ORFs, and utilization of varied ribosomal entry sites in transcript structure (Johnstone, et al., 2016; Kochetov, 2008; Lee, et al., 2012). Determining the impact of AS, ATRI/ATRT, and ATLI/ATLT, or their combinations is challenging and becomes more so with advancing organismal complexity, where, for instance, human genome has only ∼1/3rd of total exons as protein-coding (Aspden, et al., 2023). Additionally, ATRI/ATRT contributes ∼4-fold higher alternative nucleotides (nt) in transcript region than AS (Shabalina, et al., 2014), which contributes to non-trivialities in deciphering their influence on coding sequence (CDS). Complexity resulting from these AS and related processes is not limited to humans but is also noted in mouse, where splicing exon profiles from noncoding regions were recapitulated for human chr21 (Deveson, et al., 2018).

Considering the multifaceted roles of gene architecture in influencing transcriptional and translational processes, previous studies unveiled several features concerning introns and exons. These suggested a reduction in their length and increase in count was pivotal with the rise in organismal complexity (Koralewski and Krutovsky, 2011; Movassat, et al., 2019; Zhu, et al., 2009). However, literature reports have relied majorly on AS, with limited emphasis on other associated processes (ATRI/ATRT and ATLI/ATLT) and their synergistic roles in shaping proteome. Previous studies emphasizing splicing event notation prioritized defining them through pairwise exonic/transcript comparisons. For instance, the numeric symbolic notation is employed by Foissac et al. and Sammeth et al. (Foissac and Sammeth, 2007; Sammeth, et al., 2008), to identify AS events and subtypes and splice graph-based data structures have been employed by Xing et al., and Vaquero-Garcia et al., (Vaquero-Garcia, et al., 2016; Xing, et al., 2006) to uncover similarity and complex events in transcriptome data. Modifications of associated approaches, specifically in the splice graphs, have been developed to uncover the splicing complexity (Vaquero-Garcia, et al., 2016), discover unique exon junctions, and increase computational efficiency in transcriptomic analyses (Sterne-Weiler, et al., 2018). Though such approaches are robust in efficiently elucidating splicing events of interest, they do not always distinguish between CDS/UTR and their intervening regions, the importance of which has been greatly emphasized in literature (Shabalina, et al., 2014). Highlighting this research gap, Reixachs-Solé et al., (Reixachs-Solé and Eyras, 2022) also proposed a systematic inferable integration of transcriptome and proteome data. Cumulatively, challenges in exploring the extent of AS and related processes impact on proteome diversity necessitates a tool/methodology to analyze mRNA-mediated effects on the translated proteome through systematic accommodation of the intron-exon definitions in the CDS region.

To circumvent the challenges hindering the elucidation of transcriptional and translational process’s impact on gene’s protein diversity, we focused on tracking exonic loci throughout gene architecture genomic coordinates, which is usually not considered in pairwise splicing event analysis. We developed a systematic exon translational feature-centric framework, Exon Nomenclature and Classification of Transcripts (ENACT), to streamline computational tracking of exonic events while facilitating manual and automatic interpretation of their features. These include splice site variations, coding/noncoding exon property, and their combinations with exonic loci incorporated through genomic and coding genomic coordinates. Thus, ENACT provides ways to assess proteome impact of exon variations (including indels) from transcriptomic and translational processes, especially inadequacies promulgated by AS, ATRI/ATRT, and ATLI/ATLT.

## Materials and methods

### Overview of ENACT framework

#### A) Unique indexing of exon to construct gene architecture and define alternate/constitutive features to exon

1. <u>RISO selection:</u> For a given gene, we select an isoform having the maximum number of coding exons from a set of curated isoforms (NCBI RefSeq proteins having ‘NP_’ prefix) and define it as a Reference ISOform (RISO). If the number of coding exons is identical in two or more isoforms, then the one with the longest length is selected as RISO. If a gene has no ‘NP_’ prefixed isoforms, RISO can be chosen from all known isoforms using similar criterion (Box –I).
2. <u>Reference set of exons (*RSOEx*):</u> Initially, exons of RISO constitute the Reference Set Of Exons (*RSOEx*), which are subsequently populated with exons from other isoforms (*nrIsfSet*) based on overlap of genomic coordinates (GC) (Box-I and represented in Figure 1). This procedure of exon selection (for *RSOEx*) involves segregation of overlapping exons (*OlEx*, with GC overlap to *RSOEx*) from non-overlapping exons (*NolEx*, without GC overlap to *RSOEx*). The latter consists of singleton exons (*Nolex- A*), which are added to *RSOEx* and another subgroup of exons (*NolEx-B*) having self GC overlaps (described in routine *defineSuboverlapExons* (Box-1) and shown in Figure 1(B1)). Subsequently, *NolEx-B* subgroups are processed to identify *Qualifier_exon_* in following way: from a subgroup, an exon of minimum length of at least ten amino acids (30 nt) or the longest (if maximal exon length among overlapping entities (*GoEx*) is <30 nt) is chosen as representative exon (*Qualifier_exon_*) for *RSOEx* set (Figure 1(B1)). Others left in the above subgroup are moved to a set defined *Exon_variant_*, which also consists of *OlEx* exons. Subsequently, *RSOEx* exons are sorted on their GCs and are assigned relative position (linear index, first character in block-II, Figure 1E) (see algorithm Box -1).
3. <u>Prevalence of exon entities:</u> The next character of block-II (Figure 1E) depicts alternate/constitutive property of exons, defined by their GC consistency in transcripts. An exon is considered alternate (shown as ‘A’) if it lacks uniform presence in all transcripts and is constitutive (depicted using ‘G’) otherwise. Additionally, certain exonic positions are depicted by ‘F’, representing these as occurring in all isoforms; however, with splice site variations.

#### B) Relationship definition between *RSOEx* and *Exon_variant_*

The previous step identifies reference exon (*RSOEx*) and variant exon sets (*Exon_variant_*). In this step, we define specific splice site variant relations (5’ and/or 3’ exon) between exons, considering their GC overlap. We chose to define splice site and its variations focusing on the exonic entity (De Conti, et al., 2013), as ENACT framework involves their featurization with protein sequence.

1. <u>Splice site variability:</u> Each exonic entity in *Exon_variant_* set is compared with those in *RSOEx* and is assigned notation as follows:

a. n: It denotes different 5’ splice site (5ss) but an identical 3’ splice site (3ss) for *i^th^* entity of *Exon_variant_* to the *k^th^* entity of *RSOEx*.
b. β. c: It denotes identical 5ss but different 3ss for *i^th^* entity of *Exon_variant_* to *k^th^* entity of *RSOEx*.
c. b: It denotes different 5ss and 3ss for *i^th^* entity of *Exon_variant_* to *k^th^* entity of *RSOEx*.
d. δ. 0 (default): It denotes identical 5ss and 3ss for *i^th^* entity of *Exon_variant_* to *k^th^* entity of *RSOEx*.

**Figure 1:**
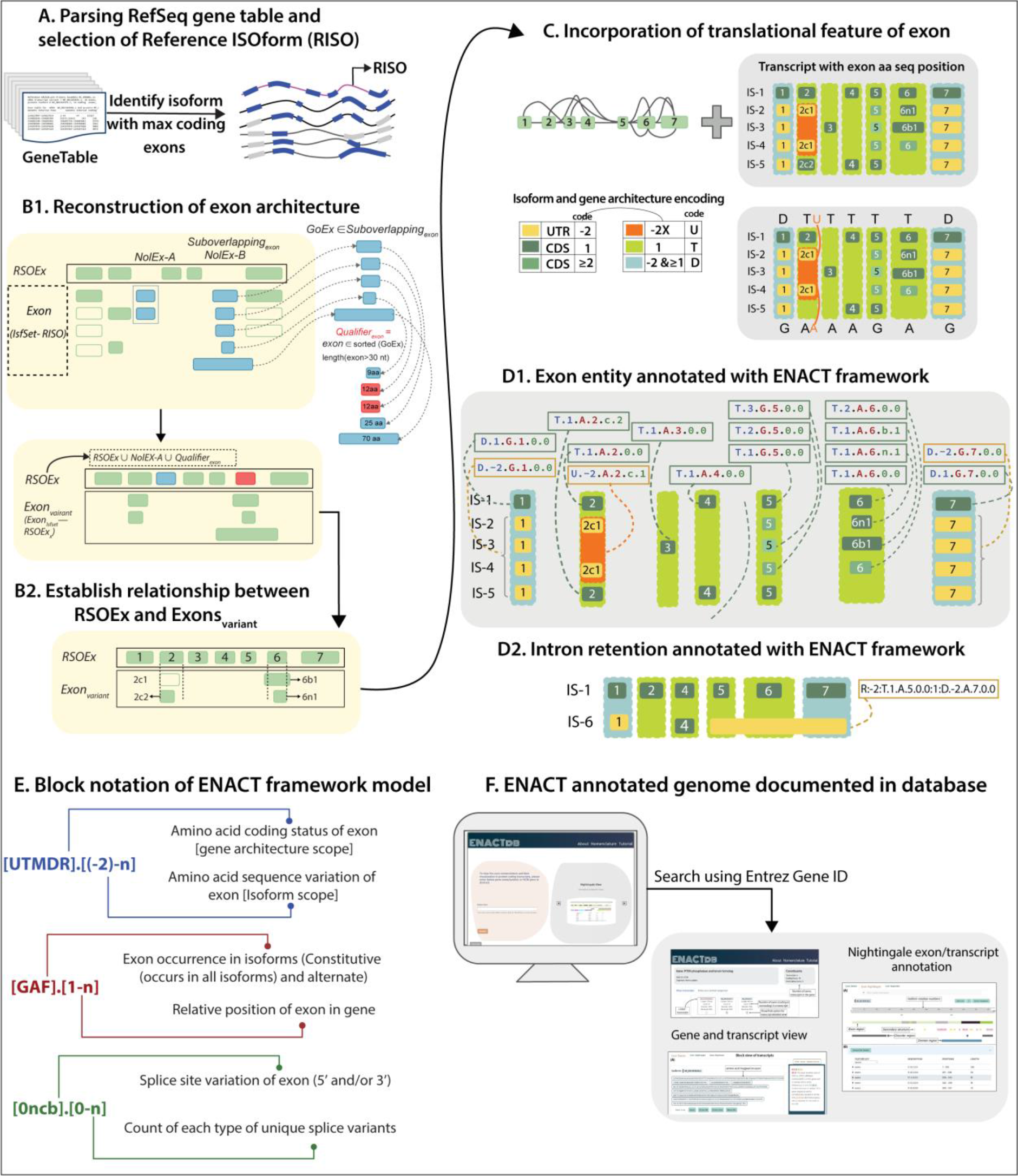
Overview of Exon Nomenclature and Classification of Transcripts (ENACT) framework. Figure illustrates main steps of the ENACT algorithm (A-D) and the framework model (E) for a gene. A) Source data of genes, comprising curated models of transcripts with GC and CGC concerning each exon is obtained from NCBI RefSeq in gene table format (Blue and grey color regions represent coding and noncoding exons, respectively). The reference isoform (RISO) is selected based on the maximum number of coding exons (see methods). B1) RISO exons constitute primary set of reference exons (*RSOEx*), which are later screened for candidate exons from other isoform: a. singleton non-overlapping (concerning GC of *RSOEx*) exons (*NolEx-A*) and b. a selected exon (Qualifier) from a group of overlapping (*NolEx-B*) exonic entities (based on their GCs) that has a minimum length of 30 nt or otherwise the maximum length. Thus, *RSOEx* set consists of non-overlapping exons (sequentially ordered based on GCs) from all isoforms. A set of *Exon_variants_* is maintained, comprising entities with GC that overlap *RSOEx*. An exon with GC overlap with ≥2 *RSOEx* instances is tagged as Intron retention and maintained separately. B2) Exons in *Exon_variant_* are compared with those of *RSOEx* to define splice site variants (alternate splice site) as *n/c/b* instances (see methods for details). C) CGCs establish translational attributes for each exon (*RSOEx*/*Exon_variant_*). The relative position (considering their GCs) of all exonic entities (*RSOEx* and *Exon_variant_*) and their prevalence in isoforms are noted as linear index and constitutive/alternate state attributes. Thus, each exon is assigned its relative position, translational feature, occurrence, and splice site variation (if known). D1) The exon attributes noted in the previous steps are combined to construct a 6-character alphanumeric notation defined as an exonic entity (EUID). D2). The intron retention instances are identified in step B and annotated with IR codes. E) The 6- character EUID is segregated into three blocks (I, II, III) of 2 characters each. G) The description and notation for each exon character are illustrated in the figure. The annotated exons of genes encoded in five model organisms are documented in (ENACTdb) database with distinct visualization views of exon annotations.

The above notations (*n/c/b*/0) are utilized as first character of block-III and represent either extension or shortening of the exon length. It should be noted that when an exonic entity (*Exon_variant_*) showed GC overlap to more than one entity in *RSOEx*, we defined them as intron retention events, which is described later (see methods section E).

1. <u>Occurrence of splice site changes:</u> The variant relationship between entities of *Exon_variant_* between *RSOEx* is facilitated by their GC overlap. To accommodate and acknowledge more than one GC overlapped entity (*Exon_variant_*) with *RSOEx* instance, we track them by counting their occurrences. For instance, in Figure 1(B2) *i^th^* and *j^th^* entities of *Exon_variant_* have identical 5ss but vary in 3ss (*c* variation) with *k^th^ RSOEx* instance, inferring its two distinct *c* variations. We note their distinct occurrence (1 for *i^th^* and 2 for *j^th^*), which are stored as 2^nd^ positional variable in block-III. Additionally, this value for *RSOEx* entity will be represented by 0 and is incremented with previously unobserved splice site variant. Interestingly, the number of *n*/*c*/*b* events of a specific exon locus (determined by block-I linear indexed attribute) in a gene can be obtained by extracting number of this position.

#### C) Amino acid coding (translational) attribute of exons defined in *RSOEx* and *Exon_variant_*

The translational attribute of the exon is shown in block-I using two characters (Figure 1E). The first character (global scope) identifies the region of gene architecture exonic locus belongs, and second character (local scope) denotes its isoform specific amino acid contribution. Global scopes are denoted by letters ‘T’, ’U’, ‘D’, ’M’, and ‘R’, and local scopes have been defined by numeric identifiers ‘-2’ to ‘n’, with details as follows:

- ‘**T**’ (**T**ranslated): Depicts exonic locus containing CGC in all its isoform instances. Hence, it is a part of CDS regime globally in gene architecture. Amino acid sequence (‘aas’) of current locus need not be uniform among its occurring isoforms and may yield different sequences, considering alternate promoter-driven ATLI in N-terminal, non-3n exon skipping driven reading frame change in the middle or truncation in C-terminal (ATLT). Accordingly, subtype local scope is necessary to encompass complete translational features and is defined as below:

`o` 1: Reference ‘aas’ contribution of an exon.
`o` ≥2: Increment counter denoting the number of different ‘aas’ variants observed for an exon with the same GCs (because of ATLI, ATLT, or reading frameshift).
`o` -1: Depicts premature stop codon in the upstream exon; hence, concerning locus lacks ‘aas’ contribution despite having CGC.
- **‘U’** (**U**n-Translated): Depicts exonic locus lacking CGC for all its isoform instances and is a part of UTR regime throughout gene architecture. The numeric tag ‘-2’ always denotes isoform-specific scopes (local) of such instances.
- **‘D’** (**D**ual): are exonic loci that are part of CDS and UTR regime, thus showing inconsistent CGC (variable and none) in its occurring isoforms. Additionally, for a locus to have a global ‘D’ scope, it should have at least two local scope instances with tags of ‘-2’ and ‘1’.
- **‘M’** (**M**icro): Depicts locus having CGC of ‘1nt’ and indicates single nucleotide protein-coding exon in the CGC region. The ‘0’ local tag marks these instances.
- **‘R’** (**R**etention): Denotes exon undergoing intron retention, described in the later section (methods E).

#### D) EUID construction

The attributes discussed in the method sections C, A (A2, and A3), and B (B1, and B2) comprise block-I, II, and III attribute groups respectively. The exonic features are encoded as alphanumeric characters joined sequentially by ‘.’ (in the order of block-I, block-II, and block-III) as demonstrated in Figure 1E, yielding a 6-character long alphanumeric string descriptor termed as Exon Unique Identifier (EUID). Briefly, the first block denotes translational attribute, and the second block indexes the relative linear position of the exon in a gene, followed by a feature of exon occurrence. The third block depicts the exon splice site variations.

The Subset of EUID characters can be used to construct sub-features by invoking EUID^k^, where ’k’ represents one or more attributes or their combinations.

#### E) Intron Retention (IR)

The intron retention events, filtered in the methods section B1, are treated as a special case of exon nomenclature where 6-character EUID is insufficient to capture details of a retention event (as it involves retention of exon/intron region between exons). We take EUIDs of two exons and combine them with 3 other identifiers separated by colon symbol (‘;’) to construct IR-EUID (Figure 1(D2)). The first identifier is the alphabet ‘R’ to recognize that the exon is involved in IR, followed by a digit describing its amino acid coding attribute (the same notation is used as described before in the methods section C). The third and fifth identifiers are exon EUIDs, between which the intron/exon region is retained to form the IR exon. The fourth identifier is a numeric character showing the number of retention events observed involving exons and their variants. We use 0 as the default value of this counter. For instance, “R:1:U.-2.A.2.n.1:0:T.1.A.3.0.0” is an IR identifier and depicts amino acid coding exon having the retention of intron region between exons U.-2.A.**2**.n.1 to T.1.A.**3**.0.0. It is the first instance involving exons 2 and 3 (shown in bold as these are the relative positions of exons in a gene, and another instance of IR is illustrated in Figure 1(D2)).

Using ENACT, we have annotated protein coding genes of five representative organisms viz. *Caenorhabditis elegans* (worm), *Drosophila melanogaster* (fruit fly), *Danio rerio* (zebrafish), *Mus musculus* (mouse), and *Homo sapiens* (human) (see supplementary text S1). These annotations are available on webserver http://www.iscbglab.in/enactdb.

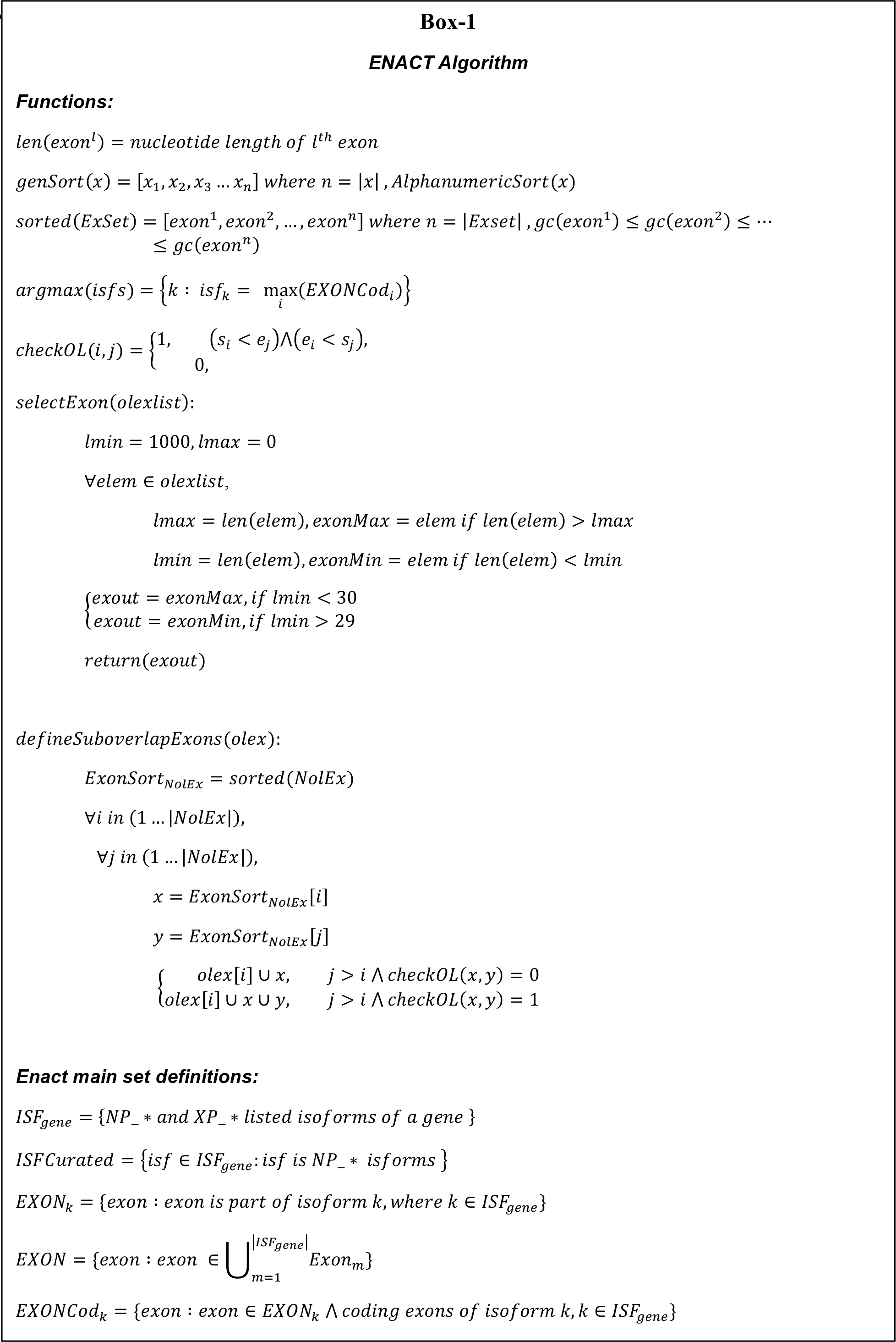

## RESULTS AND DISCUSSION

To understand the synergistic effects of transcriptomic processes and intron-exon organization within gene architecture, we constructed ENACT framework, designed to reveal and categorize features of exonic loci influenced by AS, ATRI/ATRT, and ATLI/ATLT processes. This is essential to comprehend the functional variations or translational regulation disseminated through the choice and arrangement of exons from regions of coding sequences (CDS) and untranslated regions (UTR).

### ENACT framework and exonic entity interpretation

In ENACT framework, we emphasize exonic locus as anchor position by establishing a reference set of exons (Figure 1(B1), see methods) and associate them with a linear positional index. Implementation of former is non-trivial considering alternate exon-rich mammalian transcriptomes and the splice site variability of exonic locus that presents several GC overlapping exonic polymorphs. These complexities are systematically addressed in ENACT framework (methods and Box-1). After linear indexing of exons, ENACT procedure accrues following attributes to merge them in a 6-character EUIDs (discussed in methods) from listed transcripts: a) exon translational feature, signifying potential of exonic region to code protein sequence or UTR contribution (block-I), b) exon prevalence feature, indicating alternate and constitutive nature among transcripts (block-II), and c) loci coordinate variability feature, indicating alternate splice sites for exon locus (block-III).

The combination of block-II’s linear index and block-III’s splice site variational attribute defines an exon entity instance. As we have considered exonic GCs in formulating ENACT exon entity, it is pertinent to note that every unique splice site variant (referred in block-III) of an exon (in reference to *RSOEx* instance) is a distinct instance for exonic featurization, even though its linear index is related to *RSOEx* exon instance, considering its GC overlap (described in methods, Figure 1(B2)). Therefore, block-III differing exon entities are independent of their parent *RSOEx* instance and could acquire separate or similar block-I attributes to their parent. For example in Figure 1C, exon-2’s *RSOEx* (coding (T)) and *Exon_variant_* instances (2c1 is noncoding (U)) highlight varying translational attributes in the global scope (U/T) (see attributes in methods) as they undergo CDS-UTR feature change because of splice site alteration. Alternatively, the assignment of identical ‘T’ scope for exon-6’s *‘n’* and *‘b’* splice variants highlight that global coding scope is maintained irrespective of *Exon_variant_* instances (Figure 1C). Altogether, these indicate that co-interpretation of block-II and block-III attributes, supplements relating *Exon_variant_* to *RSOEx* instances, while facilitating distinction in interpreting their translational features, the scope of which has been listed above and will also be discussed in the next section.

### Interpretation of AS, ATRI/ATRT, and ATLI/ATLT through ENACT

ENACT block featurization of exon and its attribute extraction through EUID^k^ (see methods) enables systematic assessment of intra-transcript exonic composition impacted by AS, ATRI/ATRT, and ATLI/ATLT processes.

We illustrate features of the ENACT nomenclature representation using an example of a hypothetical gene having ten exons in Figure 2. Before discussing events, we describe the exon features that can be derived from their nomenclature. Three of 10 exons are noncoding and have EUIDs assigned as U.-2.G.1.0.0, U.-2.F.9.0.0, and U.-2.A.10.0.0 (block-I global scope ‘U’, local ‘–2’). Their block-II linear position index attribute (EUID^4^) shows that U.-2.G.1.0.0 is the first constitutive entity, U.-2.F.9.0.0 is constitutive-like (EUID^3^=F code), and U.- 2.A.10.0.0 is the last and alternate exon. The second exon is assigned EUID of global scope D, indicative of its transitioning coding nature in its UTR/CDS regime, where local scope ‘1’ in TR-7 reflects its CDS participation and ‘aas’ contribution and scope ‘-2’ in other transcripts is indicative of its participation as UTR. The EUIDs of exons 3 to 8 imply them as coding exons (block-I global scope ‘T’ and local scope and ≥1), where exons 3 and 5 occur in all transcripts but show splice site variants (block-II, prevalence scope ‘F’), notated by their block-III attribute group.

**Figure 2:**
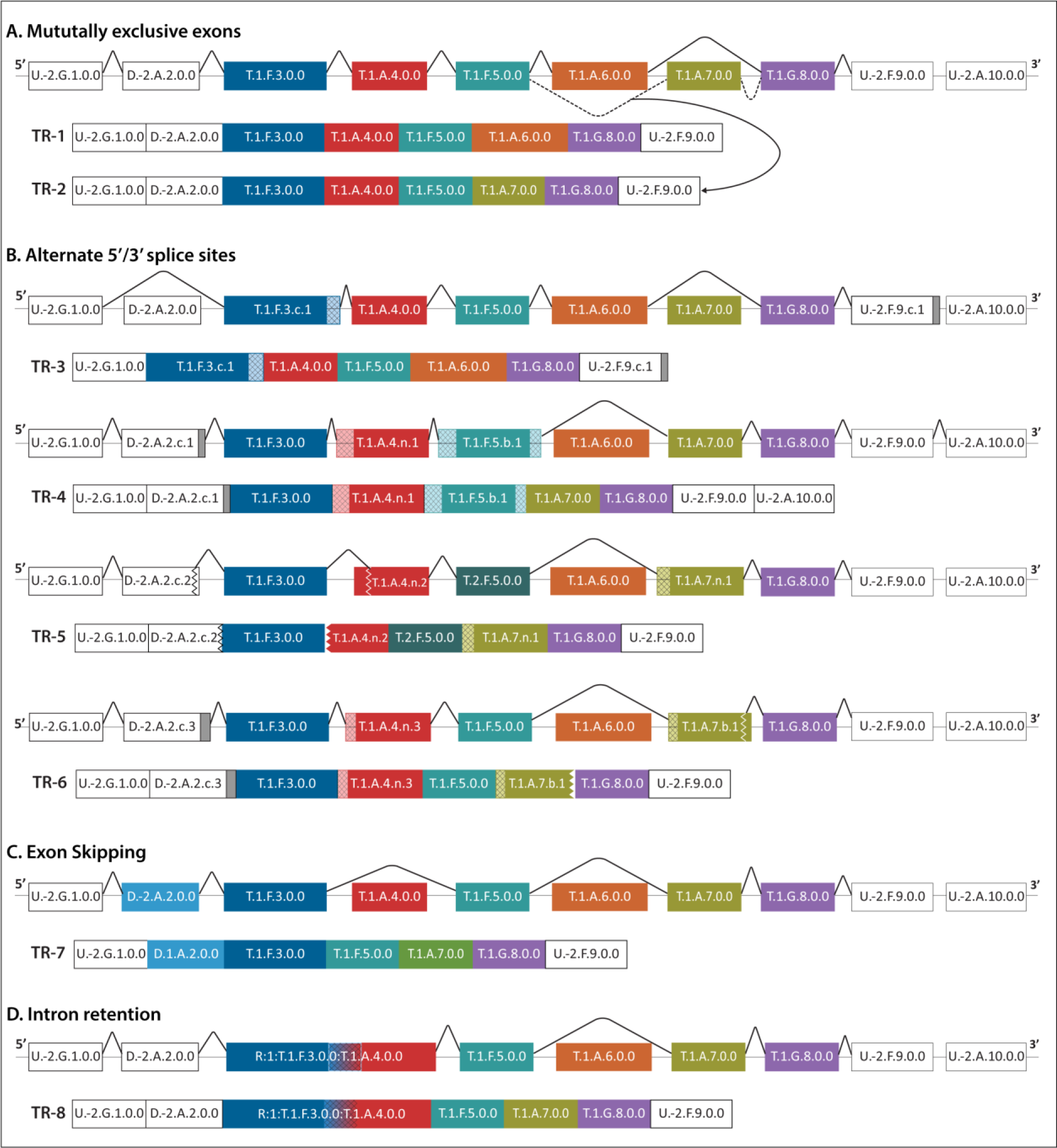
Schematic illustrating events involving Alternative Splicing (AS), Alternative Transcription (ATR), and Alternative Translation (ATL). **Figure represents a hypothetical** example of various AS, ATR, and ATL splicing events. Arrangement of exons in gene architecture representing their combination is shown followed by the resulting transcript. Exons are represented as rectangular boxes with a unique identifier (EUID) assigned to them within transcripts. Block-I feature variability of an exon is shown in variable colors. Non- coding exon is filled in white color, partial coding is filled in grey color and coding exon is filled in various colors making distinction from each other. Exon 5’ or 3’ splice site variations are represented by crosshatched filled rectangles for extensions and jagged end for shortening. Inference of AS, ATR and ATL events is facilitated by their inter-transcript comparison. ATR and ATL events involve alternation in UTR and CDS, where ATR events are inferred by alternate first or last exons and ATL events are alternate first or last coding exons or differences in block-I local scope within shared exons. AS events involving ES and A(ss) are depicted based on their block-II and block-III attributes. IR event is shown as gradation of two exon colors involved in IR. AS events, such as MXE and IR, need comparison of more than 1 loci, and their interpretation is helped by event outline before listing the transcript’s exon composition.

### Alternate transcription initiation (ATRI) and termination (ATRT) inference

ATRI and ATRT processes leave their imprint by exerting either different termini exons or exons with variable UTR lengths affecting little on AS in CDS region. Changes in 5’UTR have potential to modulate several attributes concerning further processing of nascent transcript and translational efficiency, as noted in Churbanov et al. and Resch et al. (Churbanov, et al., 2005; Resch, et al., 2009). Similar changes in 3’UTR can alter the localization of resulting product and add to possibility of distinct polyadenylation site preference as discussed in Batt et al., (Batt, et al., 1994). As shown in Figure 2, all listed transcripts share a common start exon, which is constitutive member in the UTR regime, indicating identical transcription start site; however, variable 3’UTR exons is noted. Inter-transcript comparison of TR-1 and TR-4 indicated ATRT in the latter as a consequence of alternative exon 10 participation. Additionally, exon 9 showed distinct splice site choices while being terminal when TR-1 and TR-3 are compared (Figure 2); however, they shall not be considered ATRT as splice site choices are strong indicators of splicing processes, as noted from study by Pal et al. (Pal, et al., 2011).

### Alternate translation initiation (ATLI) and termination (ATLT) inference

These phenomenon indicate differential termini coding exons in transcripts or differential translational start and termination sites. As depicted in Figure 2, exon-2 is an alternate exon entity (skipped in TR-3), whereas in TR-7 its inclusion introduces upstream ATLI when compared to other TRs. This highlights the crucial feature of differing translation start site with introduction of ‘D’ global scoped exons that transitions between UTR-CDS regimes. Similarly, as shown in Figure 1C, exons 1 and 7 are constitutive in block-II and introduce ATLI and ATLT in IS-2 to 5 compared to IS-1. ATLI/ATLT sites can also be introduced in ‘T’ global scoped exons, inferred by a change in their local coding scope values of ≥2 (2^nd^ position in EUID). However, scope of exons in introducing ATLI/ATLT shall be considered in context to their positioning near termini and interior of transcripts. The exon near termini shows association of local coding scope change with first and last coding exons became highly indicative of ATLI/ATLT, whereas in interior, they often indicate translation frame change from RISO. For example, in Figure 1D1, exon-5 harbors ‘aas’ variation (represented through shades of green color and changed local- scope characters: 1, 2, and 3), indicating ATLI from exon-5 in IS-2 and IS-4 (block-I local scope 2, Figure D1). Conversely, exon 5’s block-I local scope 3 in IS-3 is a consequence of reading frame change, when ATLI in exon-3 is employed. Similar frame changes can also be noted in *RSOEx* instances of exon-6 in IS-4.

### Alternative Splicing (AS) inference

AS modulates exonic composition in the transcript primarily in 4 major events, a) exon skipping (ES), b) mutually exclusive events (MXE), c) alternate 5’ (5ss) or/and 3’ (3ss) splice site (A(ss)) of the exon, and d) Intron retention (IR). EUIDs note ES and A(ss) as functions of its block-II and block-III features, indicative of their locus-specific inference.

Deduction of ES and A(ss) is straightforward considering the block-II (EUID^3^) and block- III (EUID^5^) features. Additionally, invoking block-III along with block-II for EUID^3^=A has the potential to identify two distinct alternate exon subtypes: case a) locus undergoing alternate prevalence with no splice variant (ES(only)), noted by identical GC’s and EUID^5^ = 0 among all instances in transcripts sharing linear positioning index (defined by block-II, EUID^4^), and case b) locus undergoing splice variants while being alternate (A(ss)), noted by GC inconsistency with EUID^5^ = *n*/*c/b* for some or all instances sharing linear positioning index. Such segregation of attributes along with similar block-II feature ‘F’ would be essential in recognizing the change in exonic composition for evolutionarily related genes, where previous studies noted distinct preference of ES and A(ss) in the species (Koralewski and Krutovsky, 2011). In coding regime of the example illustrated in Figure 2, exon loci 4, 6, and 7 are ‘T’ global scope (block-I) with alternating entities, where exon-6 lacks consistent block-III attributes, whereas exons 4 and 7 undergo ‘*n’* and ‘*b’* block-III subtype variations. Thus, implying A(only) nature of exon locus 6 and A(ss) for loci 4 and 7. Additionally, block-II’s ‘F’ prevalence scope, by definition, highlights entities to be uniformly prevalent in all known transcripts with noted variations in splice sites, highlighting additional A(ss) subtypes.

Unlike other AS events, IR and MXE events need consideration of more than exon locus and transcripts. IR is identified by the ‘R’ code of Exon in TR-8, which shows the retention of the intron region between exons 3 and 4. MXE are exon groups that do not co-occur in transcripts. Exons 6 and 7 exemplify this subtype as these two loci lack co-occurrence in known transcripts. Detection and interpretation of these events may seem straightforward; however, it becomes non-trivial if candidate exons for MXE exhibit variations in block-III (due to splice site changes), making subjected choice and consideration of MXEs to multiple exon candidates. ENACT can circumvent such issues by focusing on GCs to formulate exon entities and appropriately detect such MXEs through their relative position. The exon featurization represented in block-I and block-III provides locus-based choice to interpret MXE while acknowledging their accrued variations (splice site and/or translational attributes) rather than constructing multiple exons sets for MXEs. Similarly, IR events in the ENACT framework are depicted between exonic locus along with their block-III variations, illustrating multiple AS events that could be readily displayed in a single transcript representation.

### Functional diversity inference illustration by ENACT entities

The previous section exemplifies interpretation of ENACT entities composing transcripts. Here, we extend the significance of it in comprehending transcriptomic/proteomic diversity in isoforms introduced through a coordination of AS and/or its related processes (ATLI/ATLT). We selected two genes ADAM8 and WNK4 for case studies to demonstrate the co-interpretation of functionally induced variations and ENACT framework-based exon-centric annotation.

#### A. ADAM8

The ADAM8 gene encodes a protein belonging to the membrane-anchored disintegrin and metalloprotease proteinases family that cleaves the extracellular domain of several cell surface proteins and receptors (Fourie, et al., 2003) while being involved in various cellular functions such as inflammation, immunomodulation, neutrophil activation/mobility, immune cell migration, osteoclast stimulating factor, and neurodegeneration (Romagnoli, et al., 2014; Schlomann, et al., 2000; Yamamoto, et al., 1999). The ADAM8 domain architecture consists of an N-terminal prodomain, a catalytic metalloproteinase domain, a disintegrin domain involved in interaction with integrins, a cysteine-rich domain followed by a transmembrane region, and a C-terminal domain probably involved in protein-protein interaction through SH3 or proline-rich regions (Knolle and Owen, 2009).

ADAM8’s gene architecture comprises 23 coding and one each of noncoding and ‘dual (D)’ exons. Of the coding exons, 17 are constitutive/constitutive-like, and the rest are alternate. We analyzed three well-annotated isoforms harboring a combination of AS events (Figure 3A). The reference isoform (IS-1; NP_0011003.3) has Pep_M12B_propep, Reprolysin (metalloproteinase), Disintegrin, and ADAM_CR (cysteine-rich domain) Pfam domains lying before transmembrane region. The skipping of exon-21 is combined with 5ss of exon-22 (ES/A(ss) linear in sequence index) (IS-2: NP_001157961.1, global sequence identity from IS- 1: 79%), leads to frameshift in subsequent exons and premature termination in exon-24. These could be noted by their EUIDs: T.2.F.22.n.1, T.2.G.23.0.0, and T.2.A.24.0.0. The resulting isoform (IS-2) lacks the proline-rich region required for protein-protein interaction and is expressed in metastatic lung cancer cell lines (Knolle and Owen, 2009). The IS-3 (NP_001557962.1, global sequence identity from IS-1: 82%) exhibits skipping of exons 2 to 4, affecting the local reading frame in exons 5 to 7. Interestingly, the reading frame is restored from exon 8 onwards by the 5ss AS event (Figure 3A). Due to a change in the amino acid sequence in IS-3, the pro-domain cannot be identified in the N-terminal region of this isoform, suggesting that it may have constitutive metalloproteinase activity. Furthermore, IS-3 also lacks 2 out of 4 glycosylation sites, which are part of the pro-domain (Srinivasan, et al., 2014) but hold one of the conserved Glutamate (158E) essential for the catalytic removal of the pro- domain (Hall, et al., 2009). Further experimental studies would provide information on the enzymatic activity and biological role of IS-3.

**Figure 3:**
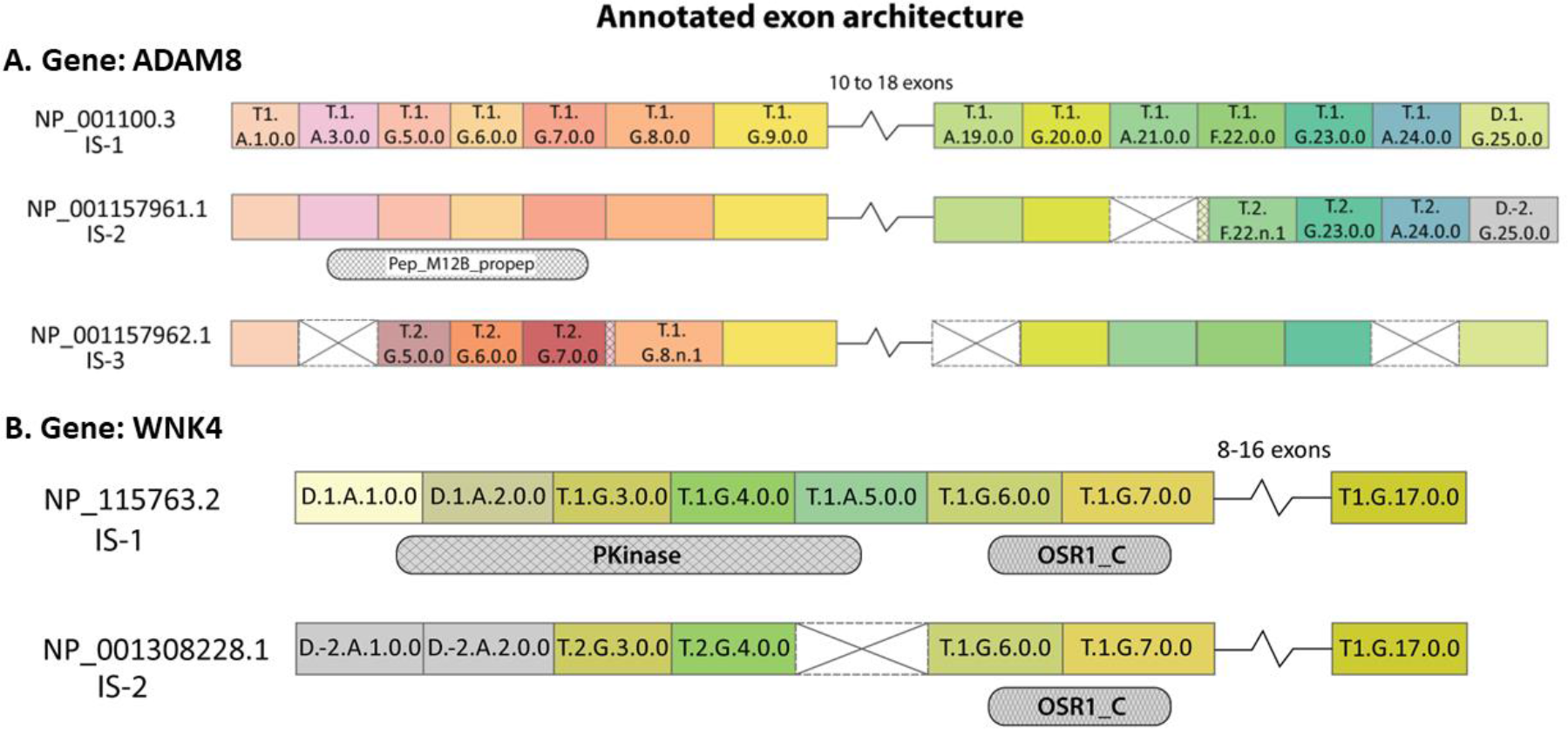
**ENACT annotation of ADAM8 and WNK4 isoforms and exons**. The NCBI gene identifier is shown for each isoform, and exons are represented as colored rectangle boxes with their respective EUID. The absence or skipped exon is shown with a crossed empty rectangle box. The small extension of exon rectangle boxes with crisscross-filled lines represents extended exon boundaries due to alternate splice sites (5ss/3ss). If an exon does not show variation with exon(s) in the isoform above, then the EUID is not labeled. The break shows that exons intervening in the region do not change in the isoforms. The jagged edges of a rectangle represent 5ss or 3ss alternate splice sites. The isoforms sharing Pfam domains for a region are shown below exon layout. In ADAM8 isoforms, IS-3 lacks the Pep_M12B_propep Pfam domain. Similarly, IS-2 of WNK4 lacks Pkinase domain in the N-terminal region.

Thus, through an integrated ENACT annotation, the exonic variation is readily localized to amino acid sequences and subsequently to functional domain region, providing inferable insights of exon variations (including indels) on the functional attributes of isoforms.

#### B. WNK4 gene

The WNK4 gene belongs to the conserved ‘With no lysine (WNK)’ group of serine/threonine kinases (STK) in eukaryotic organisms. These have been named because of their atypical positioning of catalytic lysine in subdomain II instead of I, as in other STKs. WNK4 is expressed primarily in the kidney, where WNK4 and other members of the family have a role in modulating the balance between sodium chloride re-absorption and renal potassium ion secretion (Murillo-de-Ozores, et al., 2021) by regulating the activities of cation-coupled cotransporters (SLC12, NCC), ion channels (ENaC) and ion exchangers (Moriguchi, et al., 2005; San-Cristobal, et al., 2008). The mutation in the WNK4 gene is associated with a rare genetic type of hypertension called pseudo-hypoaldosteronism type 2 (PHA2). The protein has two Pfam domains (Protein kinase and Oxidative-stress-responsive Kinase1 C-terminal domain (OSR1_C), while the rest of the sequence is intrinsically disordered. The OSR1_C encompasses the Pask-Fray 2 (PF2) domain, which is known to interact with RFX[VI] motif and suppress the activity of kinase domain (Murillo-de-Ozores, et al., 2021).

A total of 13 isoforms are listed in the NCBI RefSeq database. Of these, two isoforms are reviewed, *i.e.,* having experimental validation (Figure 3B). Among the 19 exons, the first two are ‘Dual’ as they are coding in some transcripts and non-coding in others. These two exons are encoded as coding in the reference isoform (IS-1: NP_115763.2), resulting in Pfam Protein kinase domain (Figure 3B) being assigned to the N-terminal region of the protein. In contrast, the first two exons are non-coding in IS-2 (NP_001308228.1, global sequence identity from IS-1: 66.9%) with ATLI in exon 3 (EUID: T.2.G.3.0.0), leading to a change in amino acid of exons 3-4 due to frameshift (evident from EUID^2^: T.2.G.3.0.0 and T.2.G.4.0.0). Interestingly, exon-5 ES event restores the reading frame, and the rest of protein sequence is maintained as in the reference isoform, which is also evident from the EUIDs of exons. IS-2 shows kinase domain loss but supports OSR1_C (PF2) domain. The latter domain is suggested to interact with SPAK/OSR1 protein, suggesting that IS-2 may act as a sequestering factor for them and affect their biological function.

## Discussion

In the present study, we presented an initiative to standardize exon relative position, with tracking their variations (including indels), and associate them with other annotations, allowing inference of multifaceted roles that exonic locus can encompass within gene transcripts. ENACT framework processes intra-gene transcripts to annotate exonic loci, with specific attention to their multiple GC and CGC polymorphs to accrue exon attributes. We found utility of these resulting attributes in enabling two different interpretations for gene transcripts: a) Independent interpretation of exonic loci, where block-I attribute indicates positioning of an exon in CDS/UTR or both; block-II provides linear index of loci in gene architecture and prevailing feature of loci, including its multiple polymorphs and block-III highlights polymorph specific subtype nature and relationship to linear index; b) Comparison of transcripts facilitates interpretation of ATRI/ATRT, ATLI/ATLT, and AS events. Additionally, UTR/CDS region inter–transcript variability is driven by splice variants (block-III) of coding/noncoding exonic loci (block-I attributes ‘T’, ’U’). Systematic standardization elucidated certain exons attain attribute ‘D’ (under block-I), indicative of their dual nature occurring both as CDS and UTR in transcripts. Thus, inter-transcript variability is extended by varying the coding role of exonic loci. Moreover, if dual exons are noted of alternative feature, these will hold a greater potential in inferring genes’ ability to utilize ATLI/ATLT sites.

The collective potential of ENACT annotation in inferring functional repertoire in isoforms was illustrated for genes ADAM8 and WNK4. ADAM8 gene harbors isoform diversity employing ES and A(ss) event in the N-terminal of IS-3, where the latter (exon-8, EUID^5^=*n*) rescued the frame change induced by the former (exon-3), which has compromised the identity of ‘Pep_M12B_propep’ domain. A similar phenomenon was noted in the WNK4 gene, where exons 3 and 4 underwent reading frame change in IS-2, compromising the ‘Pkinase’ domain because of ATLI but not AS; however, AS rescued the frame change induced by skipping exon- 5, leading to maintenance of ‘OSR1_C’ domain. These AS-induced rescues of frame change can be considered random or functional; however, rescue immediately after domain region indicates its functional significance to maintain similar region other than the domain.

The work of Shabalina et al. highlighted the existence of broader correlations between exon and intron-based re-arrangement in CDS, 5’ UTR, and 3’ UTRs (Shabalina, et al., 2010). Briefly, they observed a positive correlation between a) ATRI and AS in 5’UTR, b) AS in CDS region and ATRT in 3’UTR, and anticorrelation between c) AS in CDS and AS and ATRI in 5’UTR. Interestingly, the biological relevance of observations in a) and b) could be rationalized from literature (Lynch, et al., 2005; Mignone, et al., 2002; Shabalina, et al., 2010), as the former likely indicates utility of AS in splicing long UTR exons, which are selected by ATRI processes, and the latter likely indicates the purposing of distinct proteins resulting from AS to different subcellular conditions by varying alternate polyadenylation sites (PAS) (Shabalina, et al., 2010; Shyu, et al., 2008). However, the anticorrelation mentioned in (c) is challenging to explain from the current literature, which partially suggests modulation of upstream UTR without change in splicing patterns. This could be a possible mechanism to tune the tissue-specific translation rate of proteins. A careful analysis of WNK4 indicates another mechanism related to c). Here AS in CDS (exon-5 skipping) complements ATLI (exon 3), with no AS and ATRI in 5’ UTR. Since AS and ATRI in 5’UTR also have potential to influence translation and could introduce ATLI (Cenik, et al., 2010; Kramer, et al., 2013; Palaniswamy, et al., 2010; Weber, et al., 2023), in WNK4, the ATLI is driven by exon-3 compensated this role. Moreover, the ATLI followed by ES of exon-5 restores the reading frame change introduced by former (exon), resulting into partial N-terminal variability, demonstrating the impact of their combined action leading to partial diversification, and shedding light on rudimentarily understood aspect of their anticorrelation. This is speculative and requires a comprehensive study for further validation, and it could pave the way for a detailed examination to rationalize the mechanistic basis of anticorrelation (Shabalina, et al., 2010).

ENACT framework and database provide a foundation for comprehending isoform- generated diversity visually and exploring various correlations among AS and related processes on a gene-by-gene basis. This will serve as an important resource for experimentalists and computational biologists to decipher embedded functional repertoire resulting from transcriptional and translational processes.

## Acknowledgments

The authors acknowledge Deepanshi Awasthi for her critical reading of the manuscript. We also acknowledge the computing facility Param Smriti formed under National Supercomputing Mission.

## Funding

IISER Mohali

Bioinformatics Center (BT/PR40419/BTIS/137/36/2022), Department of Biotechnology under the Ministry of Science and Technology, Govt. of India

National Network Project (BT/PR40198/BTIS/137/56/2023), Department of Biotechnology under the Ministry of Science and Technology, Govt. of India

