## supplementary file for "Exon Nomenclature and Classification of Transcripts (ENACT): Systematic framework to annotate exon attributes"

**Keywords:** Alternative splicing, exon nomenclature, exon inclusion, intron retention

**S1: Overview of ENACT Database resource.**

Using our nomenclature, we have annotated exons of five widely studied model organisms, *viz. Caenorhabditis elegans, Drosophila melanogaster, Danio rerio, Mus musculus,* and *Homo sapiens* and documented them in the ENACT resource database (enactdb). Database is publicly available at http://www.iscbglab.in/enactdb/. The Table S1 summarizes the number of annotated exons/transcripts of genes encoded in five organisms available in enactdb.

**Table S1: Summary statistics of gene/transcript/exon in five model organisms.**

| Organism | Number of protein coding genes | Number of transcripts | Number of exons |
|---|---|---|---|
| *C. elegans* (Ce) | 19,972 | 28,534 | 1,25,054 |
| *D. melanogaster* (Dm) | 13,972 | 30,755 | 65,958 |
| *D. rerio* (Dr*)* | 26,374 | 48,821 | 2,68,035 |
| *M. musculus* (Mm) | 22,134 | 92,400 | 2,32,520 |
| *H. sapiens* (Hs) | 20,443 | 1,30,739 | 2,41,910 |
